# A Central Limit Theorem for Punctuated Equilibrium

**DOI:** 10.1101/039867

**Authors:** K. Bartoszek

## Abstract

Current evolutionary biology models usually assume that a phenotype undergoes gradual change. This is in stark contrast to biological intuition, which indicates that change can also be punctuated-the phenotype can jump. Such a jump can especially occur at speciation, i.e. dramatic change occurs that drives the species apart. Here we derive a Central Limit Theorem for punctuated equilibrium. We show that, if adaptation is fast, for weak convergence to hold, dramatic change has to be a rare event.

AMS subject classification: 60F05, 60J70, 60J85, 62P10, 92B99

## 1 Introduction

A long–standing debate in evolutionary biology is whether changes take place at times of speciation (punctuated equilibrium Eldredge and Gould [16], Gould and Eldredge [20]) or gradually over time (phyletic gradualism, see references in Eldredge and Gould [16]). Phyletic gradualism is in line with Darwin’s original envisioning of evolution (Eldredge and Gould [16]). On the other hand, the theory of punctuated equilibrium was an answer to what fossil data was indicating (Eldredge and Gould [16], Gould and Eldredge [19, 20]). A complete unbroken fossil series was rarely observed, rather distinct forms separated by long periods of stability (Eldredge and Gould [16]). Darwin saw “the fossil record more as an embarrassment than as an aid to his theory” (Eldredge and Gould [16]) in the discussions with Falconer at the birth of the theory of evolution. Evolution with jumps was proposed under the name “quantum evolution” (Simpson [32]) to the scientific community. However, only later (Eldredge and Gould [16]) was punctuated equilibrium re–introduced into contemporary mainstream evolutionary theory. Mathematical modelling of punctuated evolution on phylogenetic trees seems to be still in its infancy (but see Bokma [8, 10, 11], Mattila and Bokma [23], Mooers and Schluter [25], Mooers et al. [26]). The main reason is that we do not seem to have sufficient understanding of the stochastic properties of these models. An attempt was made in this direction (Bartoszek [5]) — to derive the tips’ mean, variance, covariance and interspecies correlation for a branching Ornstein–Uhlenbeck (OU) process with jumps at speciation, alongside a way of quantitatively assessing the effect of both types of evolution.

Combining jumps with an OU process is attractive from a biological point of view. It is consistent with the original motivation behind punctuated equilibrium. At branching, dramatic events occur that drive species apart. But then stasis between these jumps does not mean that no change takes place, rather that during it “fluctuations of little or no accumulated consequence” occur (Gould and Eldredge [20]). The OU process fits into this idea because if the adaptation rate is large enough, then the process reaches stationarity very quickly and oscillates around the optimal state. This then can be interpreted as stasis between the jumps — the small fluctuations. Mayr [24] supports this sort of reasoning by hypothesizing that “The further removed in time a species from the original speciation event that originated it, the more its genotype will have become stabilized and the more it is likely to resist change.”

In this work we build up on previous results (Bartoszek [5], Bartoszek and Sagitov [6]) and study in detail the asymptotic behaviour of the average of the tip values of a branching OU process with jumps at speciation points evolving on a pure birth tree. To the best of our knowledge the work here is the first to analytically consider the effect of jumps on a branching OU process in a phylogenetic context. It is possible that some of the results could be special subcases of general results on branching Markov processes (Ren et al. [29, 30]). However, these studies use a very heavy functional analysis apparatus, which unlike the direct one here, is completely inaccessible to the applied reader. We can observe the same competition between the tree’s speciation and OU’s adaptation (drift) rates, resulting in a phase transition when the latter is half the former (the same as in the no jumps case Adamczak and Miłoś [1, 2], Bartoszek and Sagitov [6]). We show here that if large jumps are rare enough, then the contemporary sample mean will be asymptotically normally distributed. Otherwise the weak limit can be characterized as a “normal distribution with a random variance”. Such probabilistic characterizations are important as there is a lack of tools for punctuated phylogenetic models. This is partially due to an uncertainty of what is estimable, especially whether the contribution of gradual and punctuated change may be disentangled (but Bokma [11] indicates that they should be distinguishable). Large sample size distributional approximations will allow for choosing seeds for numerical maximum likelihood procedures and sanity checks if the results of numerical procedures make sense. Most importantly the understanding of the limiting behaviour of evolutionary models with jumps will allow us to see the effects of these jumps, especially how much do they push the populations out of stationarity.

In Section 2 we introduce the considered probabilistic model. Then in Section 3 we present the main results. Section 4 is devoted to a series of technical convergence lemmata that characterize the speed of decay of the effect of jumps on the variance and covariance of tip species. Finally in Section 5 we calculate the first two moments of a scaled sample average, introduce a submartingale related to the model and put this together with the previous convergence lemmata to prove the Central Limit Theorems of this paper.

## 2 A model for punctuated stabilizing selection

### 2.1 Phenotype model

Stochastic differential equations (SDEs) are today the standard language to model continuous traits evolving on a phylogenetic tree. The general framework is that of a diffusion process

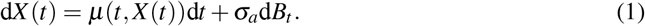

The trait follows Eq. (1) along each branch of the tree (with possibly branch specific parameters). At speciation times this process divides into two processes evolving independently from that point. The full generality of Eq. (1) is not implemented in contemporary phylogenetic comparative methods (PCMs). Currently they are focused on the OU processes

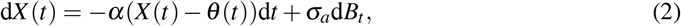

where *θ* (*t*) can be piecewise linear, with different values assigned to different regimes (see e.g. Bartoszek et al. [7], Butler and King [12], Hansen [21]). There have been a few attempts to go beyond the diffusion framework into Lévy process, including Laplace motion, (Bartoszek [4, 5], Duchen et al. [14], Landis et al. [22]) and jumps at speciation points (Bartoszek [5], Bokma [9, 10]). We follow in the spirit of the latter and consider that just after a branching point, with a probability *cp*, independently on each daughter lineage, a jump can occur. We assume that the jump random variable is normally distributed with mean 0 and variance 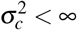. In other words, if at time *t* there is a speciation event, then just after it, independently for each daughter lineage, the trait process *X* (*t*^+^) will be

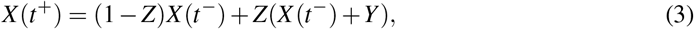

where *X* (*t^−/^*^+^) means the value of *X* (*t*) respectively just before and after time *t*, *Z* is a binary random variable with probability *p* of being 1 (i.e. jump occurs) and 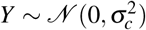. The parameters *p* and 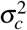 can, in particular, differ between speciation events.

### 2.2 The branching phenotype

In this work we consider a fundamental model of phylogenetic tree growth — the conditioned on number of tip species pure birth process. We first introduce some notation, illustrated in Fig. 1 (see also Bartoszek [5], Bartoszek and Sagitov [6], Sagitov and Bartoszek [31]). We consider a tree that has *n* tip species. Let *U* ^(*n*)^ be the tree height, *τ*^(*n*)^ the time from today (backwards) to the coalescent of a pair of randomly chosen tip species, 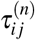 the time to coalescent of tips *i*, *j*, ϒ^(*n*)^ the number of speciation events on a random lineage, *υ*^(*n*)^ the number of common speciation events for a random pair of tips bar the splitting them event and 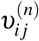 the number of common speciation events for a tips *i*, *j* bar the splitting them event. Furthermore let *T_k_* be the time between speciation events *k* and *k* + 1, *p_k_* and 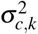 be respectively the probability and variance of the jump just after the *k*–th speciation event on each daughter lineage.

**Figure 1:**
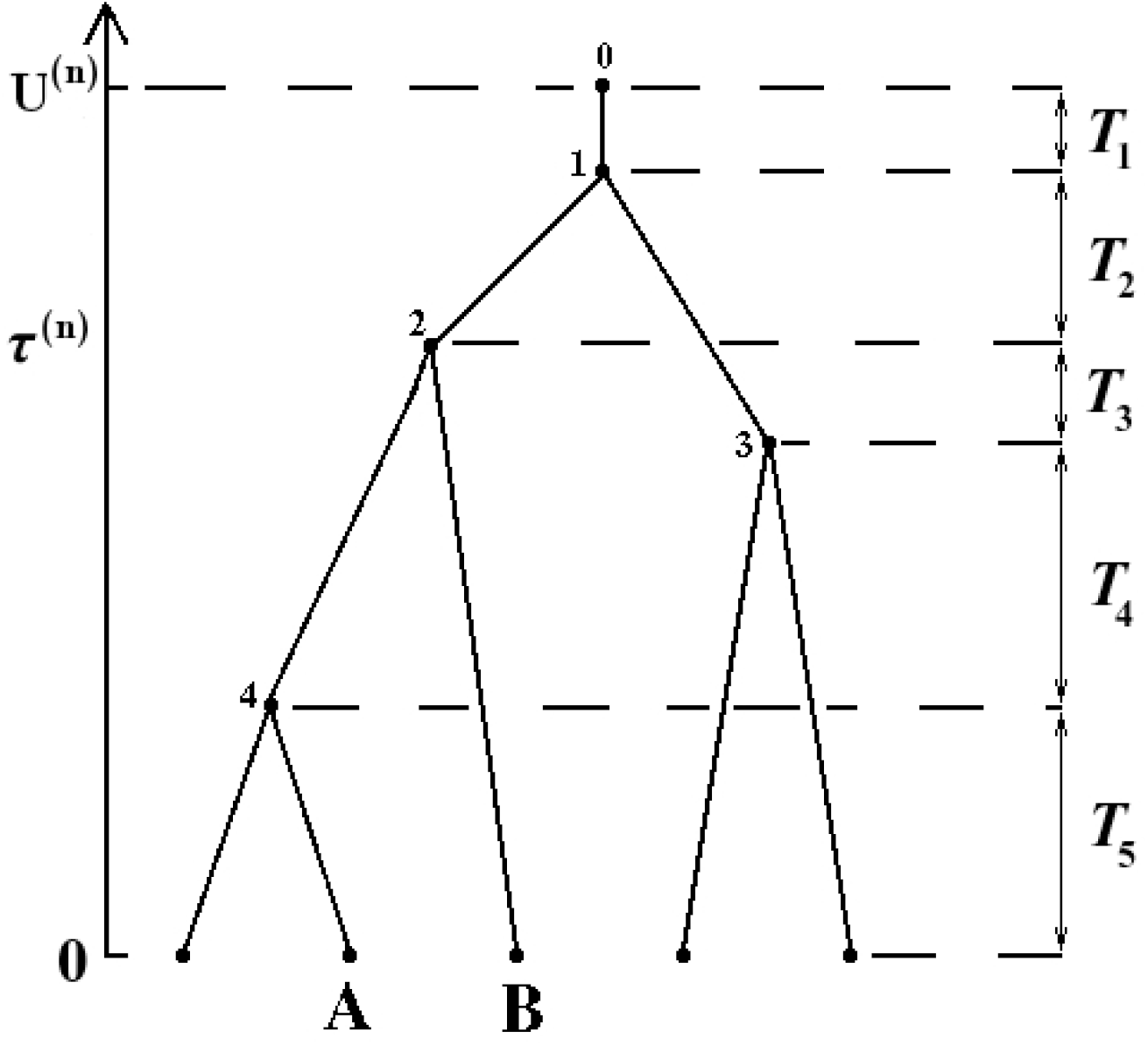
A pure–birth tree with the various time components marked on it. If we “randomly sample” node “A”, then ϒ^(*n*)^ = 3 and the indexes of the speciation events on this random lineage are 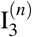 = 4 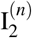 and 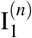. Notice that 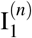 = 1 always. The between speciation times on this lineage are *T*_1_, *T*_2_, *T*_3_ + *T*_4_ and *T*_5_. If we “randomly sample” the pair of extant species “A” and “B”, then *υ*^(*n*)^ = 1 and the two nodes coalesced at time *τ*^(*n*)^. The random index of their joint speciation event is ^Ĩ^_1_ = 1. See also Bartoszek [5]’s Fig. A.8. for a more detailed discussion on relevant notation. The internal node labellings 0–4 are marked on the tree.

The above intuitive descriptions can be made more precise. We first introduce two separate labellings for the tip and internal nodes of the tree. Let the origin of the tree have label “0”. Next we label from “1” to “*n* − 1” the internal nodes of the tree in their temporal order of appearance. The root is “1”, the node corresponding to the second speciation event is “2” and so on. We label the tips of the tree from “1” to “*n*” in an arbitrary fashion. This double usage of the numbers “1” to “*n* − 1” does not generate any confusion as it will always be clear whether one refers to a tip or internal node.

#### Definition 2.1

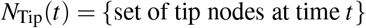

#### Definition 2.2

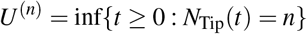

#### Definition 2.3

*For i ∈ N*_Tip_(*U* ^(*n*)^),

ϒ^(*i,n*)^ = {number of nodes on the path from the root (internal node 1, including it) to tip node *i*}

#### Definition 2.4

*For i ∈ N*_Tip_(*U* ^(*n*)^),

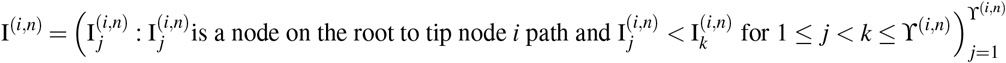

#### Definition 2.5

*For i ∈ N*_Tip_ (*U* ^(*n*)^) *and r* ∈ {1, ⋯, *n* − 1}, *let* 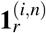 *be a binary random variable such that*

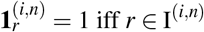

#### Definition 2.6

*For i ∈ N*_Tip_(*U* ^(*n*)^) *and r* ∈ {1, ⋯, ϒ^(*n*)^}, *let J*^(*i,n*)^ *be a binary random variable equalling* 1 *iff a jump took place just after the r–th speciation event in the sequence* I^(*i,n*)^.

#### Definition 2.7

*For i ∈ N*_Tip_(*U* ^(*n*)^) *and r* ∈ {1, ⋯, *n* − 1}, *let* 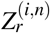 *be a binary random variable equalling* 1 *iff* 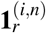 = 1 *and* 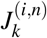 = 1, *where* 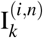 = *r*.

#### Definition 2.8

*For i, j ∈ N*_Tip_(*U* ^(*n*)^),

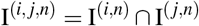

#### Definition 2.9

*For i, j ∈ N*_Tip_(*U* ^(*n*)^),

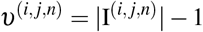

#### Remark 2.10

*We have the −*1 *in the above definition of υ*^(*i, j,n*)^ *as we are interested in counting the speciation events that could have a jump common to both lineages. As the jump occurs after a speciation event, the jumps connected to the coalescent node of tip nodes i and j cannot affect both of these tips*.

#### Definition 2.11

*For i, j ∈ N*_Tip_(*U* ^(*n*)^) *and r* ∈ {1, ⋯, max*{I*^(*i, j,n*)^} − 1}, *let* **1***^(i, j,n^) be a binary random variable such that*

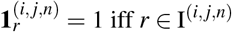

#### Definition 2.12

*For i, j ∈ N*_Tip_(*U* ^(*n*)^),

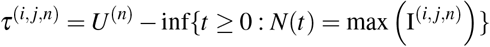

#### Definition 2.13

*For i, j ∈ N*_Tip_ (*U* ^(*n*)^) *and* 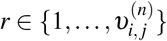 *let* 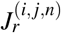 *be a binary random variable equalling* 1 *iff* 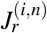 = 1 *and* 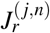 = 1.

#### Definition 2.14

*For i, j ∈ N*_Tip_ (*U* ^(*n*)^) *and r* ∈ {1, ⋯, *n* − 1}, *let* 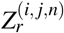 *be a binary random variable equalling* 1 *iff* 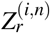 *and* 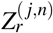.

#### Definition 2.15

*Let R be uniformly distributed on* {1, ⋯, *n*} *and* (*R, K*) *be uniformly distributed on the set of ordered pairs drawn from* {1, ⋯, *n} (i.e*. Prob((*R, K*) = (*r, k*)) = 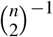, *for* 1 *≤ r < k ≤ n)*

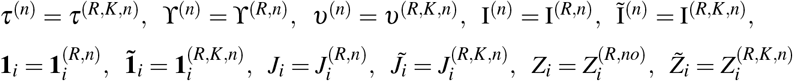

#### Remark 2.16

*For the sequences* I^(*n*)^, I^(*r,n*)^, I^(*R,n*)^, Ĩ^(*n*)^, I^(*r,k,n*)^, I^(*R,K,n*)^ *the i–th element is naturally indicated as* 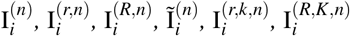 *respectively*.

#### Remark 2.17

*We drop the n in the superscript for the random variables* **1**_*i*_, 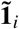, J_i_, 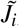, Z_i_ and 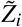 *as their distribution does not depend on n. In fact, in principle, there is no need to distinguish between the version with and without the tilde. However, such a distinction will make it more clear to what one is referring to in the derivations of this work*.

The following simple, yet very powerful, Lemma comes from the uniformity of the choice of pair to coalesce at the *i*–th speciation event in the backward description of the Yule process. The full proof can be found in Bartoszek [5] on p. 45.

#### Lemma 2.18

*For all i* ∈ {1, ⋯, *n* − 1}

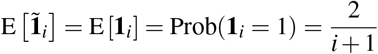

We called the model a conditioned one. By conditioning we consider stopping the tree growth just before the *n* + 1 species occurs, or just before the *n*–th speciation event. Therefore, the tree’s height *U* ^(*n*)^ is a random stopping time. The asymptotics considered in this work are when *n* → ∞.

The key model parameter describing the tree component is *λ*, the birth rate. At the start, the process starts with a single particle and then splits with rate *λ*. Its descendants behave in the same manner. Without loss generality we take *λ* = 1, as this is equivalent to rescaling time.

In the context of phylogenetic methods this branching process has been intensively studied (e.g. Bartoszek and Sagitov [6], Crawford and Suchard [13], Edwards [15], Gernhard [17, 18], Mulder and Crawford [27], Sagitov and Bartoszek [31], Steel and McKenzie [35]), hence here we will just describe its key property. The time between speciation events *k* and *k* + 1 is exponential with parameter *k*. This is immediate from the memoryless property of the process and the distribution of the minimum of exponential random variables. From this we obtain some important properties of the process. Let *H_n_* = 1 + 1*/*2 + … + 1*/n* be the *n*–th harmonic number, *x >* 0 and then their expectations and Laplace transforms are (Bartoszek and Sagitov [6], Sagitov and Bartoszek [31])

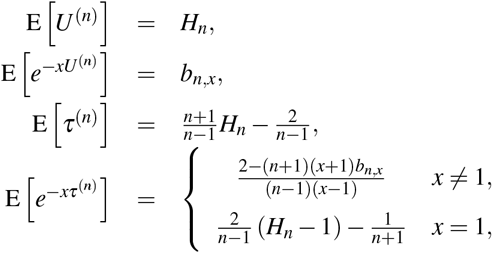

where

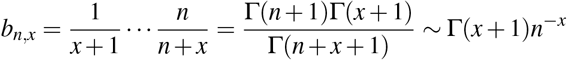

Γ(*·*) being the gamma function.

Now let *𝒴_n_* be the *σ*–algebra that contains information on the Yule tree and jump pattern. Bartoszek [5] previously studied the branching OU with jumps model and it was shown (but, therein for constant *p_k_* and 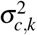 and therefore there was no need to condition on the jump pattern) that, conditional on the tree height and number of tip species the mean and variance of an arbitrary tip species, 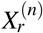, are

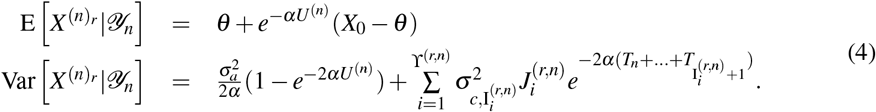

A key difference that the phylogeny brings in is that the tip measurements are correlated through the tree structure. One can easily show that conditional on *𝒴_n_*, the covariance between an arbitrary pair of extant traits, 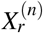 and 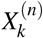 is

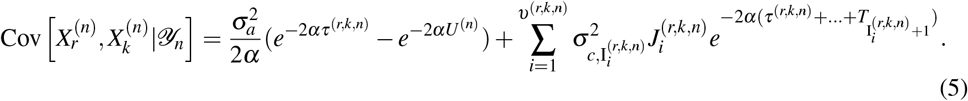

We will call, the considered model the Yule–Ornstein–Uhlenbeck with jumps (YOUj) process.

#### Remark 2.19

*Keeping the parameter θ constant on the tree is not as simplifying as it might seem. Varying θ models have been considered since the introduction of the OU process to phylogenetic methods (Hansen [21]). However, it can very often happen that the θ parameter is constant over whole clades, as these species share a common optimum. Therefore, understanding the model’s behaviour with a constant θ is a crucial first step. Furthermore if constant θ clades are far enough apart one could think of them as independent samples and attempt to construct a test (based on normality of the species’ averages) if jumps have a significant effect (compare Thms. 3.1 and 3.5)*.

### 2.3 Martingale formulation

Our main aim is to study the asymptotic behaviour of the sample average and it actually turns out to be easier to work with scaled trait values, 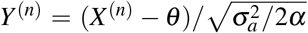. Denoting 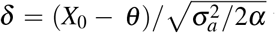 we have

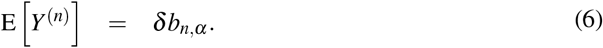

The initial condition of course will be *Y*_0_ = *δ*_0_. Just as was done by Bartoszek and Sagitov [6] we may construct a martingale related to the average

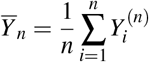

Then (cf. Lemma 10 of Bartoszek and Sagitov [6]), we define

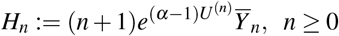

This is a martingale with respect to *F_n_*, the *σ*–algebra containing information on the Yule *n*–tree and the phenotype’s evolution.

## 3 Asymptotic regimes — main results

Branching Ornstein–Uhlenbeck models commonly have three asymptotic regimes (Adamczak and Miłoś [1, 2], Ané et al. [3], Bartoszek [5], Bartoszek and Sagitov [6], Ren et al. [29, 30]). The dependency between the adaptation rate *α* and branching rate *λ* = 1 governs in which regime the process is. If *α >* 1*/*2, then the contemporary sample is similar to an i.i.d. sample, in the critical case, *α* = 1*/*2, we can, after appropriate rescaling, still recover the “near” i.i.d. behaviour and if 0 *< α <* 1*/*2, then the process has “long memory” (“local correlations dominate over the OU’s ergodic properties”, Adamczak and Miłoś [1, 2]). In the OU process with jumps setup the same asymptotic regimes can be observed, even though Adamczak and Miłoś [1, 2], Ren et al. [29, 30]) assume that the tree is observed at a given time point, *t*, with *n_t_* being random. In what follows here, the constant *C* may change between (in)equalities. It may in particular depend on *α*. We illustrate the below Theorems in Fig. 2.

**Figure 2:**
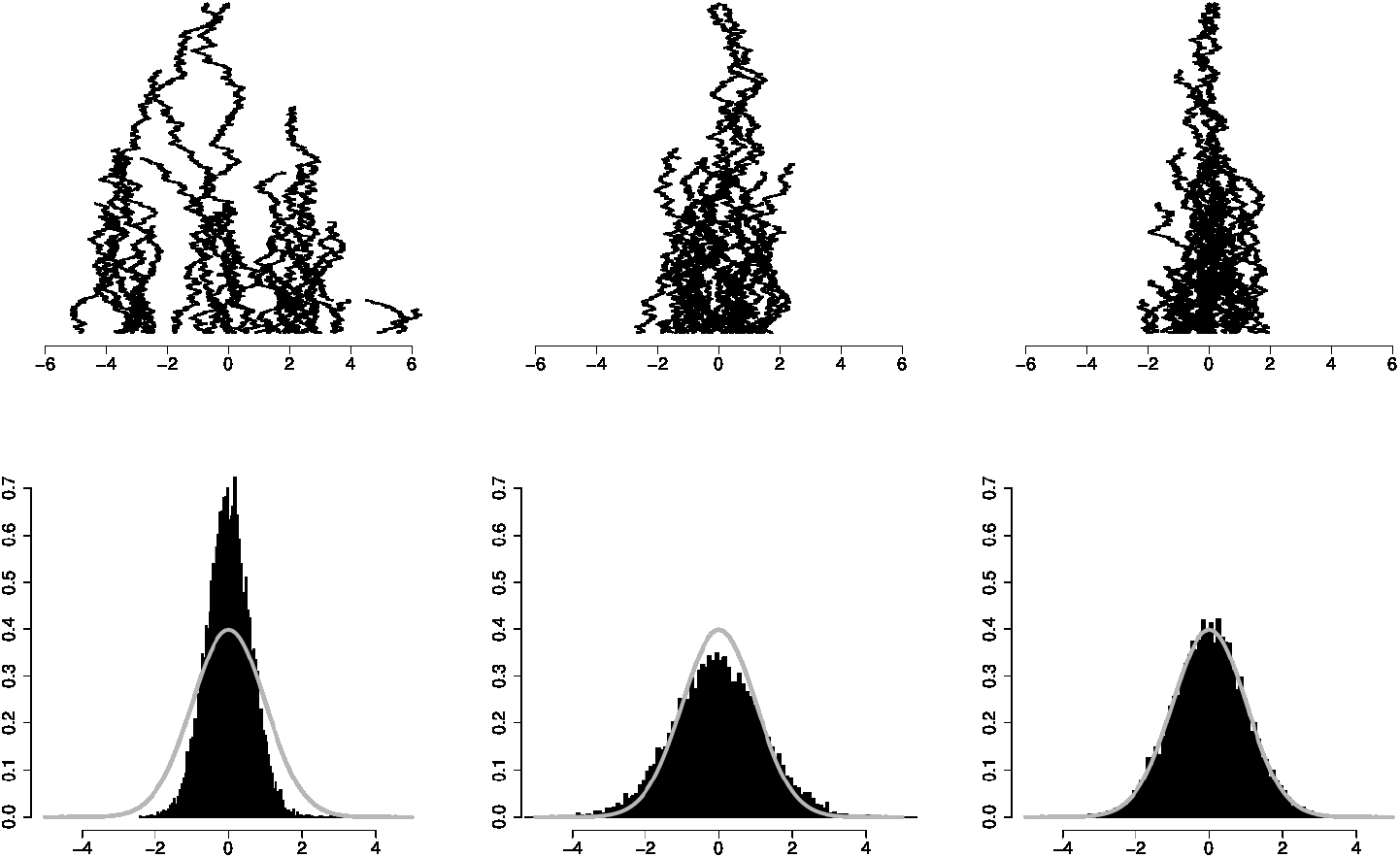
Left: *α* = 0.25 centre: *α* = 0.5 and right: *α* = 1. Top row: examples of simulated YOUj process trajectories, bottom row: histograms of sample averages, left: scaled by 1*/*Γ(3*/*2), centre: scaled by 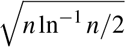, right: scaled by 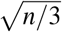. In all three cases, *p* = 0.5, 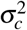, 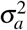, *X*_0_ = *θ* = 0. The phylogenetic trees are pure birth trees with *λ* = 1 conditioned on number of tips, *n* = 30 for the trajectory plots and *n* = 200 for the histograms. The histograms are based on 10000 simulated trees. The sample mean and variances of the scaled data in the histograms are left: (0.006, 0.360), centre: (0.033, 1.481) and right: (0.004, 1.008). The gray curve painted on the histograms is the standard normal distribution. The phylogenies are simulated by the TreeSim R package (Stadler [33, 34]) and simulations of phenotypic evolution and trajectory plots are done by newly implemented functions of the, available on CRAN, mvSLOUCH R package (Bartoszek et al. [7]).

### Theorem 3.1

*Assume that the jump probabilities and jump variances are constant equalling p and* 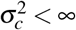 *respectively. Let* 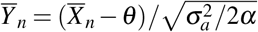 *be the normalized sample mean of the YOUj process with Y̅*_0_ = *δ*_0_. *The process Y̅_n_ has the following, depending on α, asymptotic with n behaviour*.

I. *If* 0.5 *< α, then the conditional variance of the scaled sample mean* 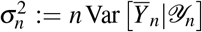 *converges in* ℙ *to a random variable* 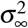 *with mean*

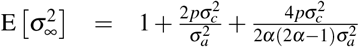 *The scaled sample mean*, 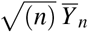 *converges weakly to random variable whose characteristic function can be expressed in terms of the Laplace transform of* 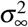

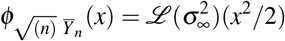
II. *If* 0.5 = *α, then the conditional variance of the scaled sample mean* 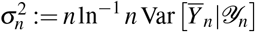 *converges in* ℙ *to a random variable* 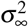 *with mean*

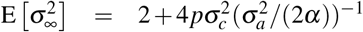 *The scaled sample mean*, 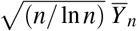 *converges weakly to random variable whose characteristic function can be expressed in terms of the Laplace transform of* 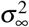

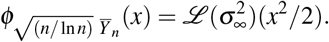
III. *If* 0 *< α <* 0.5, *then n^α^Y_n_ converges almost surely and in L*^2^ *to a random variable Y_α,δ_ with first two moments*

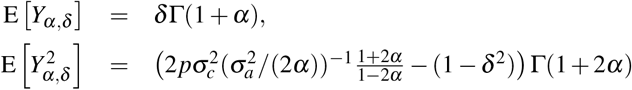

### Remark 3.2

*For the a.s. and L*^2^ *convergence to hold in Part (III), it suffices that the sequence of jump variances is bounded. Of course, the first two moments will differ if the jump variance is not constant*.

### Definition 3.3

*A subset E ⊂ 𝒩 of positive integers is said to have density* 0 *(e.g. Petersen [28]) if*

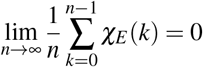

*where χ_E_* (*·*) *is the indicator function of the set E*.

### Definition 3.4

*A sequence a_n_ converges to* 0 *with density* 1 *if there exists a subset E ⊂ 𝒩 of density* 0 *such that*

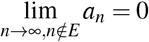

### Theorem 3.5

*If 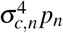 is bounded and goes to* 0 *with density* 1, *then depending on α the process Y_n_ has the following asymptotic with n behaviour*.

I. *If* 0.5 *< α, then* 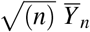 *is asymptotically normally distributed with mean* 0 *and variance* (2*α* + 1)*/*(2*α* − 1).
II. *If* 0.5 = *α, then* 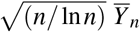 *is asymptotically normally distributed with mean* 0 *and variance* 2.

### Remark 3.6

*The assumption* 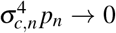 *with density* 1 *is an essential one for the limit to be a normal distribution, when α* ≥ 0.5. *This is visible from the proof of Lemma 4.5. In fact, this is the key difference that the jumps bring in — if they occur too often (or with too large magnitude), then they will disrupt the weak convergence*.

*One natural way is to keep 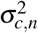 constant and allow p_n_* → 0, *the chance of jumping becomes smaller relative to the number of species. Alternatively 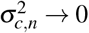, which could mean that with more and more species — smaller and smaller jumps occur at speciation. Actually, this could be biologically more realistic — as there are more and more species, then there is more and more competition and smaller and smaller differences in phenotype drive the species apart. Specialization occurs and tinier and tinier niches are filled*.

## 4 Key convergence lemmata

We will now prove a series of technical lemmata describing the asymptotics of driving components of the considered YOUj process.

### Lemma 4.1

*(Lemma* 11 *of Bartoszek and Sagitov [6])*

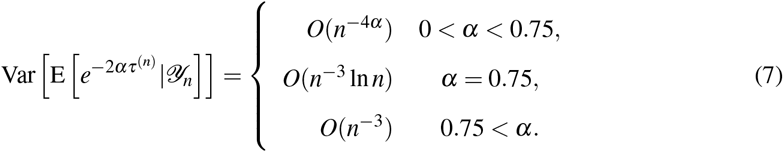

### PROOF

For a given realization of the Yule *n*-tree we denote by 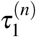 and 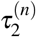 two independent versions of *τ*^(*n*)^ corresponding to two independent choices of pairs of tips out of *n* available. We have,

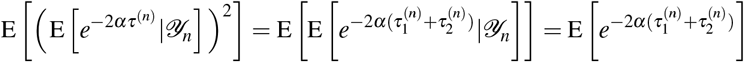

Let *π_n,k_* be the probability that two randomly chosen tips coalesced at the *k*–th speciation event. We know that (cf. Bartoszek and Sagitov [6], Steel and McKenzie [35])

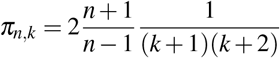

Writing

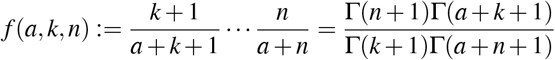

and as the times between speciation events are independent and exponentially distributed we obtain

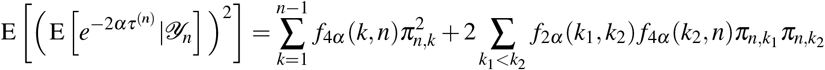

On the other hand,

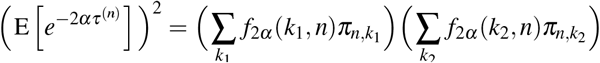

Taking the difference between the last two expressions we find

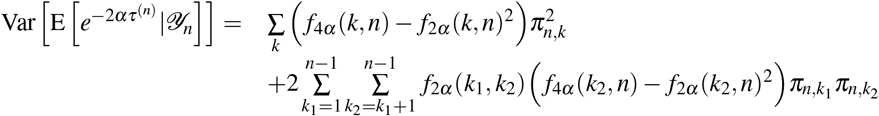

Using the simple equality

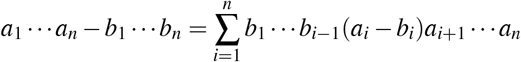

we see that it suffices to study the asymptotics of,

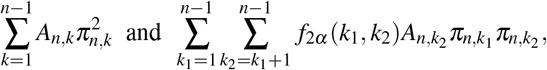

where

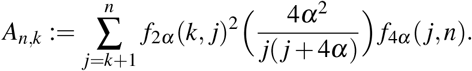

To consider these two asymptotic relations we observe that for large *n*

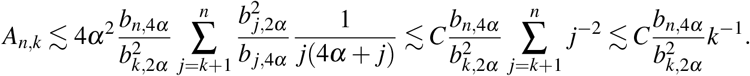

Now since 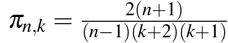, it follows

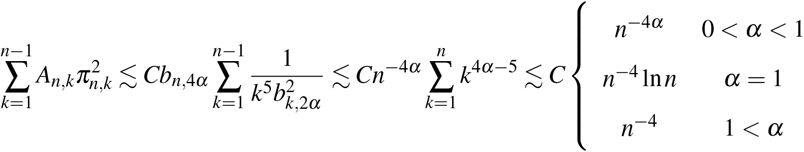

and

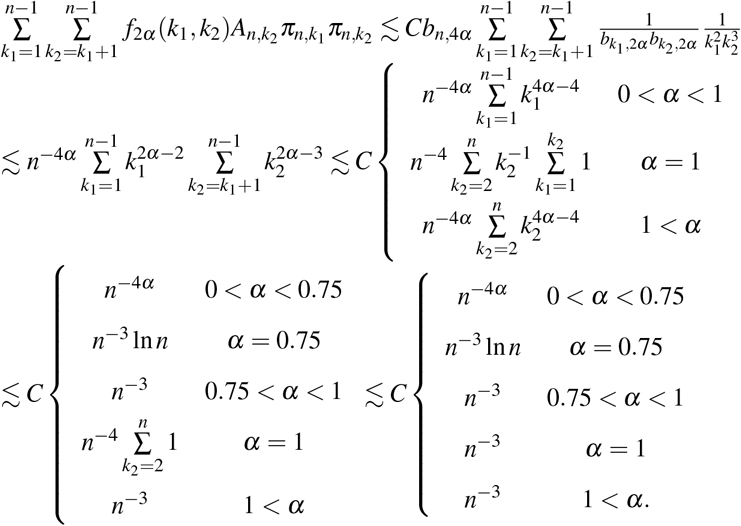

Summarizing

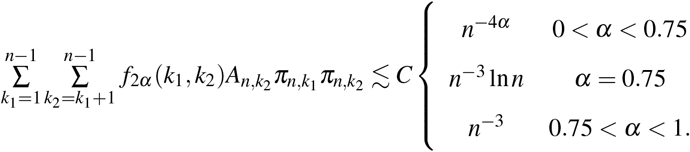

Notice that obviously 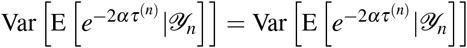.

### Remark 4.2

*The above Lemma 4.1 is a corrected version of Bartoszek and Sagitov [6]’s Lemma* 11. *There it is wrongly stated that* Var [E [*e^−^*^2*ατ*(*n*)^ *|𝒴_n_*]] = *O*(*n^−^*^3^) *for all α >* 0. *From the above we can see that this holds only for α >* 3*/*4. *This does not however change Bartoszek and Sagitov [6]’s main results. If one inspects the proof of Theorem* 1 *therein, then one can see that for α >* 0.5 *it is required that* Var [E [*e^−^*^2*ατ*(*n*)^ *|𝒴_n_*]] = *O*(*n^−^*^(2+*ε*)^), *where ε >* 0. *This by Lemma 4.1 holds. Bartoszek and Sagitov [6]’s Thm*. 2 *does not depend on the rate of convergence, only that* Var [E [*e^−^*^2*ατ*(*n*)^ *|𝒴_n_*]] → 0 *with n. This remains true, just with a different rate*.

### Lemma 4.3

*For random variables* 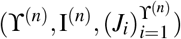 *derived from the same random lineage and a fixed jump probability p we have*

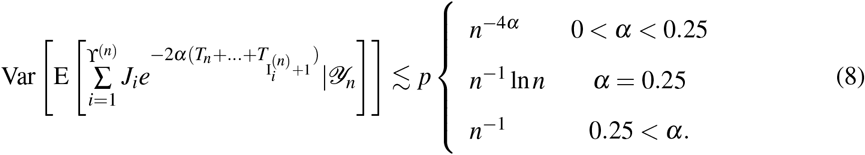

### PROOF

We introduce the random variables

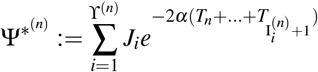

and

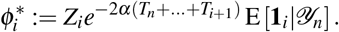

Obviously

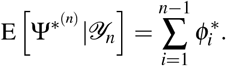

Immediately (for *i < j*)

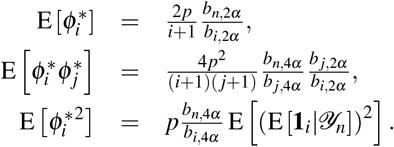

The term E [(E [**1**_*i*_|𝒴_*n*_])^2^] can be [see Lemma 11 in 6] expressed as 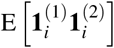 where 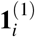 and 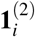 are two independent copies of **1**_*i*_, i.e. we sample two lineages and ask if the *i*–th speciation event is on both of them. This will occur if these lineages coalesced at a speciation event *k ≥ i*. Therefore

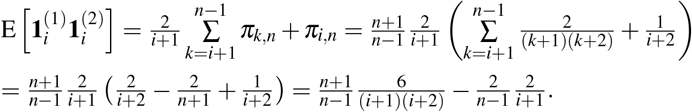

Together with the above

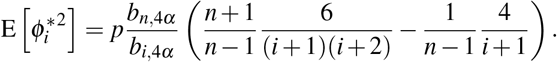

Now

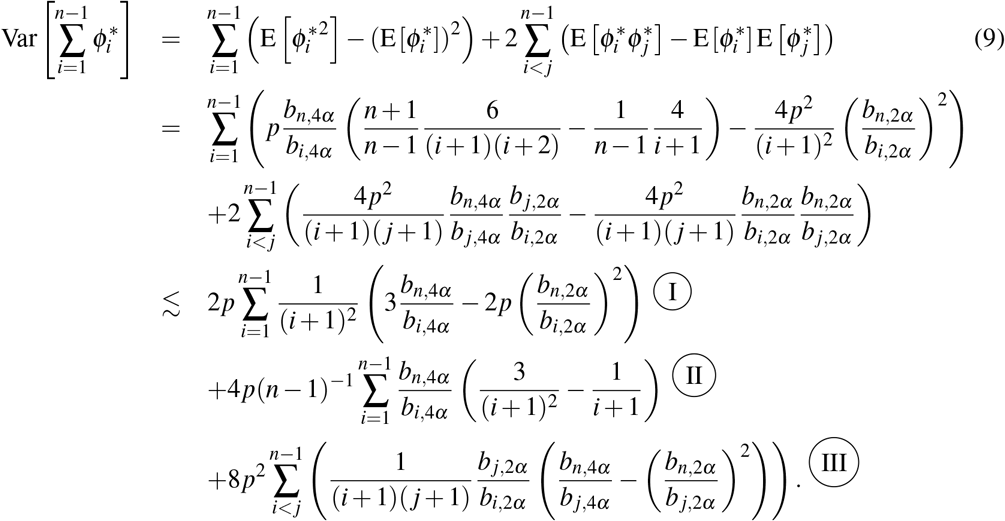

We use the equality [cf. Lemma 11 in 6]

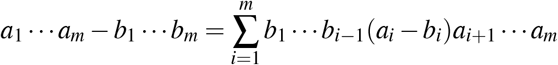

and consider the three parts in turn.

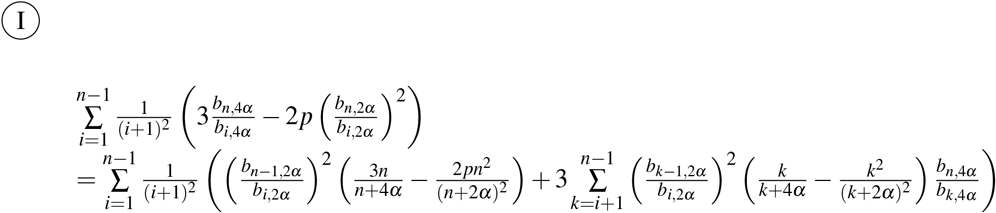

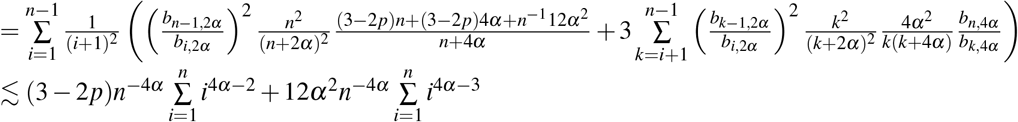

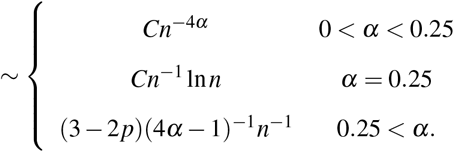

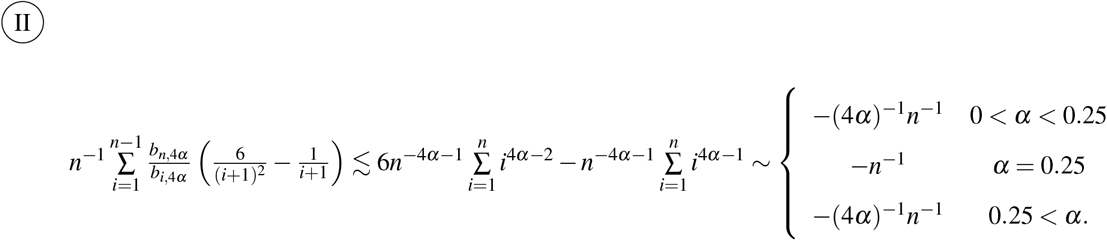

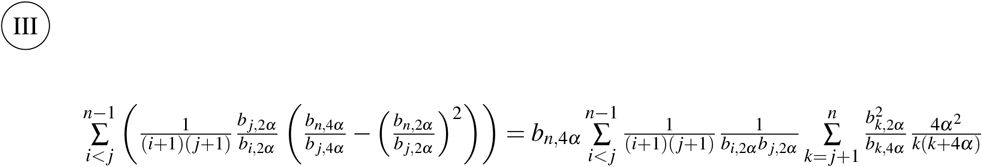

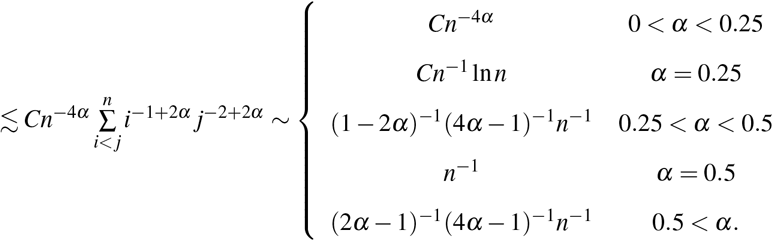

Putting these together we obtain

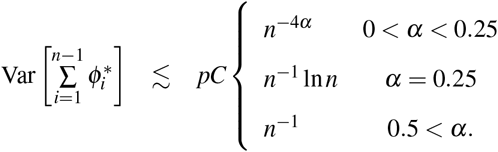

On the other hand the variance is bounded from below by 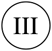. Its asymptotic behaviour is tight as the calculations there are accurate up to a constant (independent of *p*). This is further illustrated by graphs in Fig. 3.

**Figure 3:**
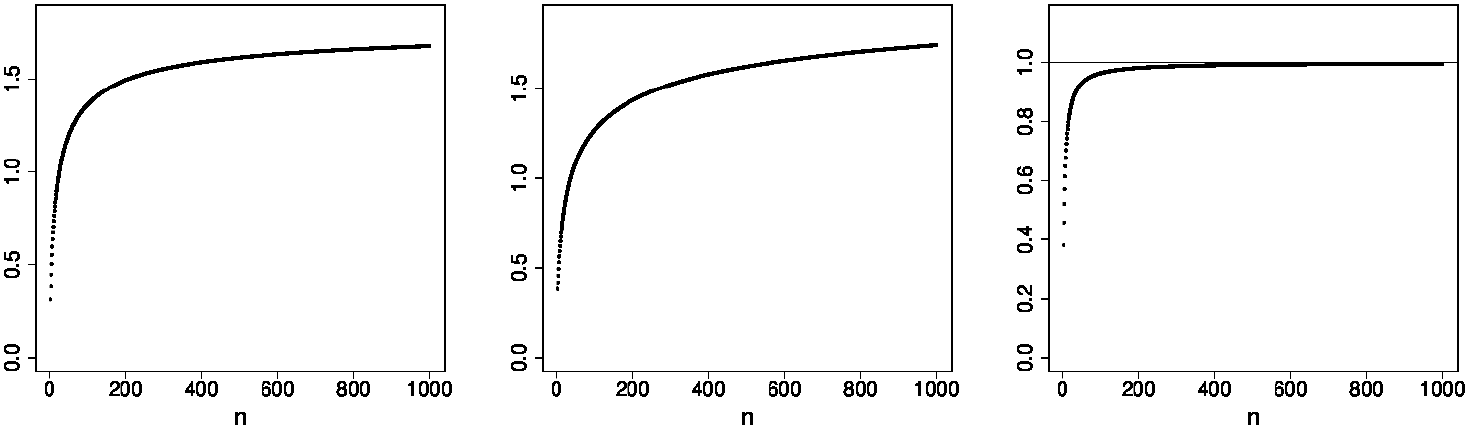
Numerical evaluation of scaled Eq. (9) for different values of *α*. The scaling for left: *α* = 0.1 equals *n^−^*^4*α*^, centre: *α* = 0.25 equals *n^−^*^1^ log *n* and right *α* = 1 equals (2*p*(3−2*p*)*/*(4*α*−1)−4*p/*(4*α*) + 32*p*^2^*α*^2^(1*/*(8*α*^2^) + 1*/*(2*α*(2*α*−1))−1*/*(4*α*^2^)^−1^−1*/*((2*α*−1)(4*α*−1))))*n^−^*^1^. In all cases, *p* = 0.5.

### Corollary 4.4

*Let p_n_ and 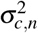 be respectively the jump probability and variance at the n–th speciation event, such that the sequence 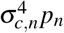 is bounded. We have*

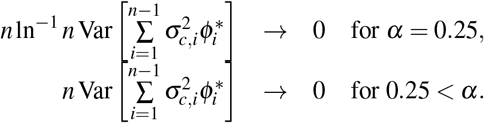

*iff* 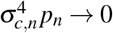 *with density* 1.

### PROOF

We consider the case, *α >* 0.25. Notice that in the proof of Lemma 4.3 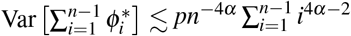. If the jump probability and variance are not constant, but as in the Corollary, then 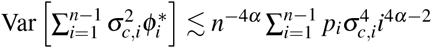.

The Corollary is a consequence of a more general ergodic property that if *u >* 0 and the sequence *a_i_* → 0 with density 1, then

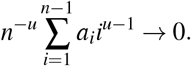

For 1 *≤ u* the above is a direct consequence of Petersen [28]’s Lemma 6.2 (p. 65). For 0 *< u <* 1 take a subsequence *N*_0_(*n*) *≤ n^u^*, such that *N*_0_(*n*) → ∞, *N*_0_(*n*)*n^−u^* → 0 and *a_N_*_0(*n*)_ *↘* 0. Then

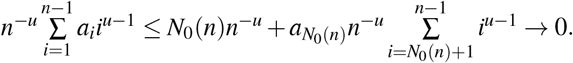

On the other hand if *a_i_* does not go to 0 with density 1, then 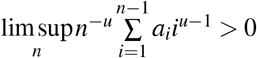.

When *α* = 0.25 we obtain the Corollary using the same ergodic argumentation for 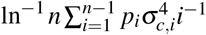.

### Lemma 4.5

*For random variables 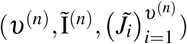 derived from the same random pair of lineages and a fixed jump probability p*

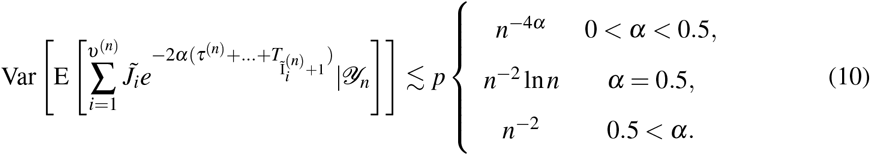

### PROOF

We introduce the notation

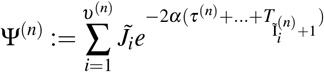

and by definition we have

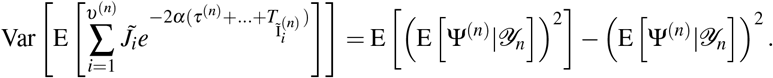

We introduce the random variable

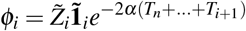

and obviously (for *i*_1_ *< i*_2_)

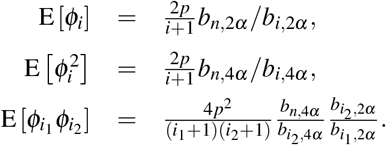

As usual let 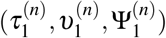 and 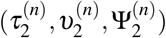 be two independent copies of (*τ*^(*n*)^, *υ*^(*n*)^, Ψ^(*n*)^) [cf. Lemma 11 in 6] and now

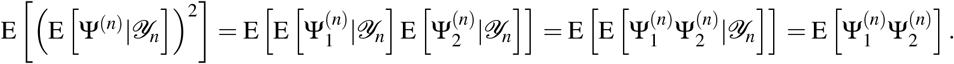

Writing out

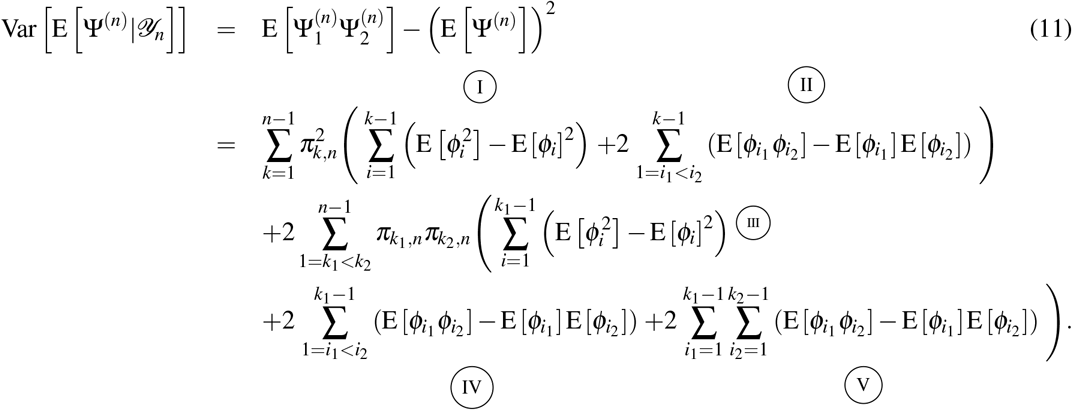

We first observe

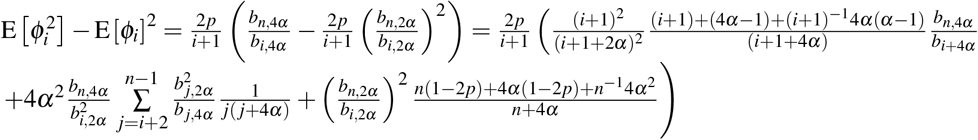

and

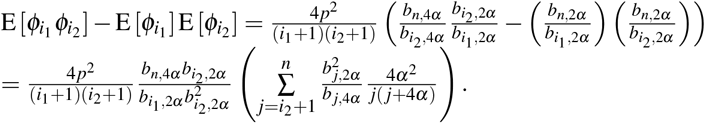

Using the above we consider each of the five components in this sum separately.

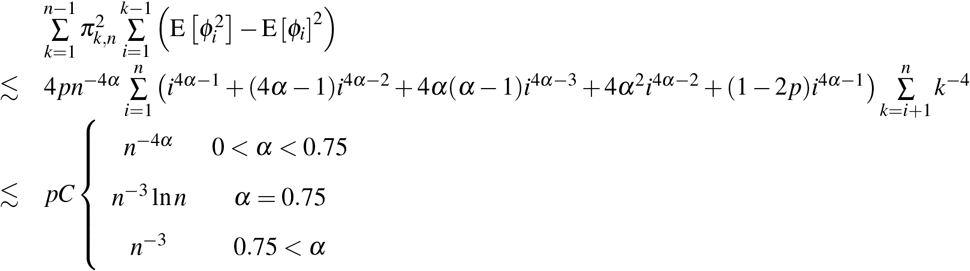

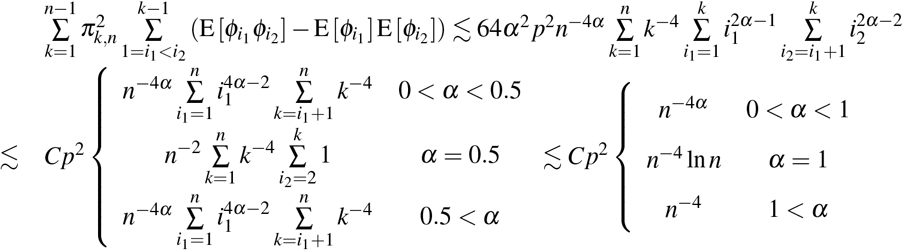

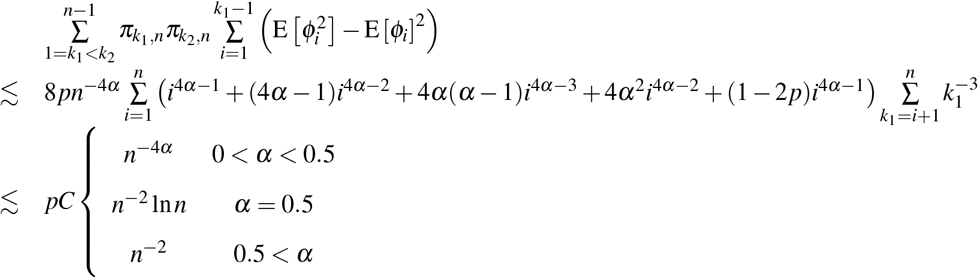

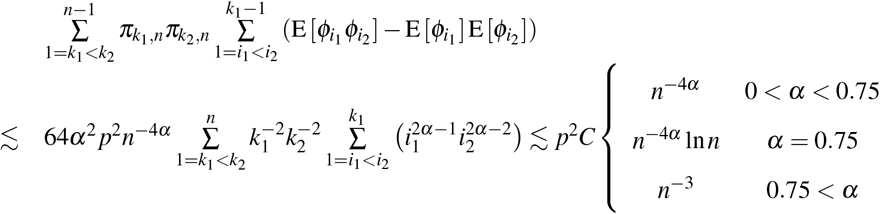

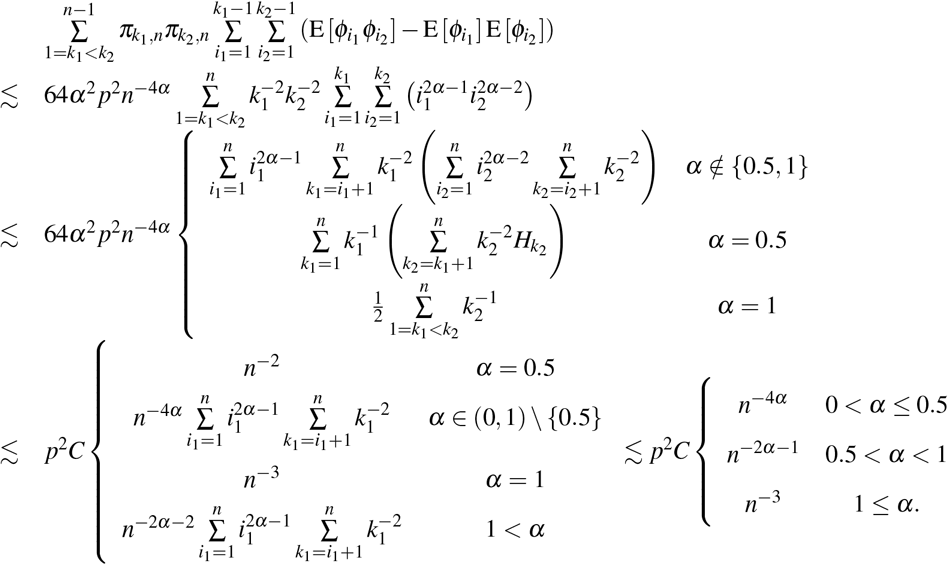

Putting I–V together we obtain

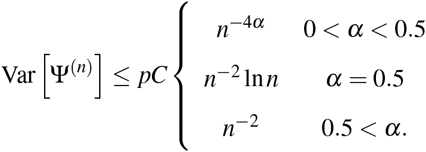

The variance is bounded from below by 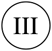 and as these derivations are correct up to a constant (independent of *p*) the variance behaves as above. This is further illustrated by graphs in Fig. 4.

**Figure 4:**
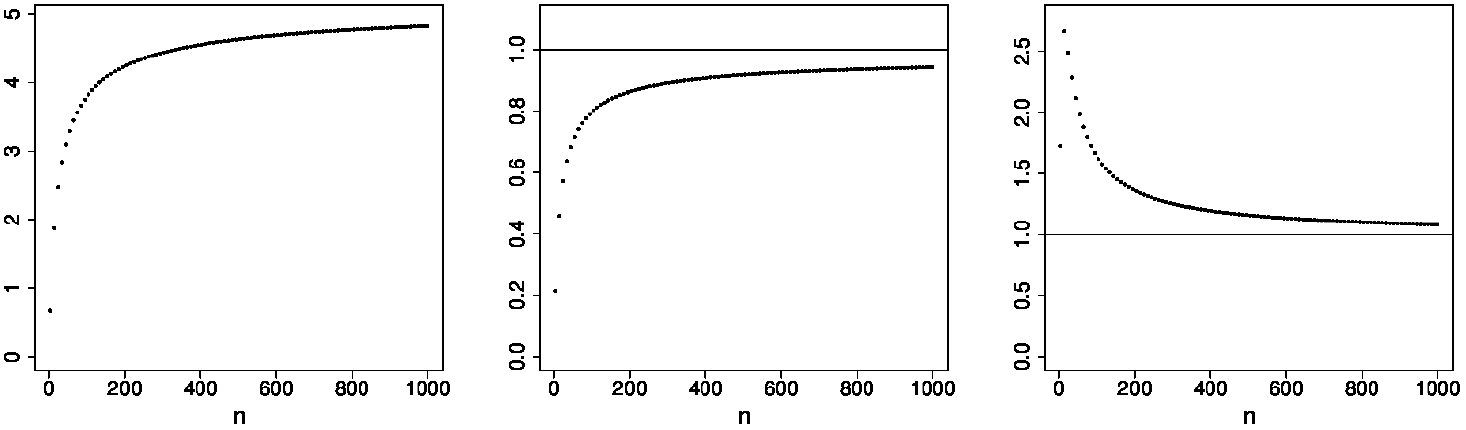
Numerical evaluation of scaled Eq. (11) for different values of *α*. The scaling for left: *α* = 0.35 equals *n^−^*^4*α*^, centre: *α* = 0.5 equals 16*p*(1 *− p*)*n^−^*^2^ log *n* and right *α* = 1 equals (32*p*(1 *− p*)(1*/*(4*α* − 2) − 1*/*(4*α* − 1))*/*(4*α*))*n^−^*^2^. In all cases, *p* = 0.5.

The proof of the next Corollary, 4.6, is exactly the same as of Corollary 4.4.

### Corollary 4.6

*Let p_n_ and* 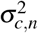 *be respectively the jump probability and variance at the n–th speciation event, such that the sequence 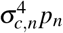 is bounded. We have*

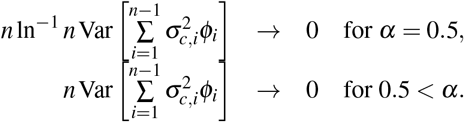

*iff* 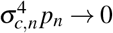 *with density* 1.

## 5 Proof of the Central Limit Theorems 3.1 and 3.5

To avoid unnecessary notation it will be always assumed that under a given summation sign the random variables 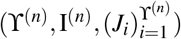 are derived from the same random lineage and also 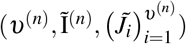 are derived from the same random pair of lineages

### Lemma 5.1

*Conditional on Y_n_ the first two moments of the scaled sample average are*

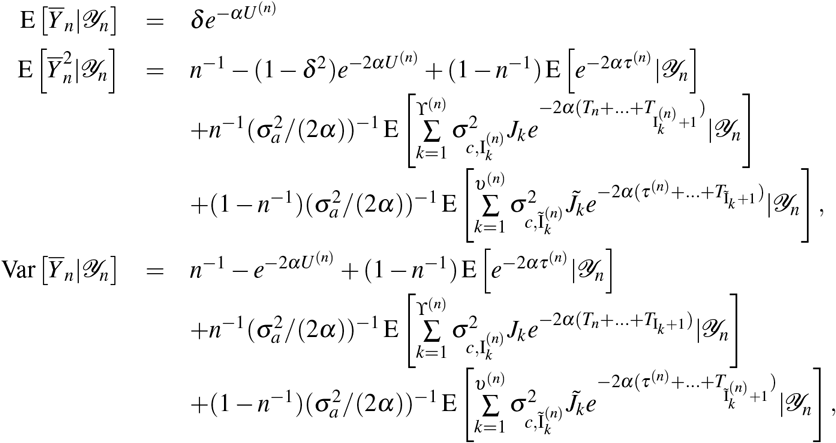

### PROOF

The first equality is immediate. The variance follows from

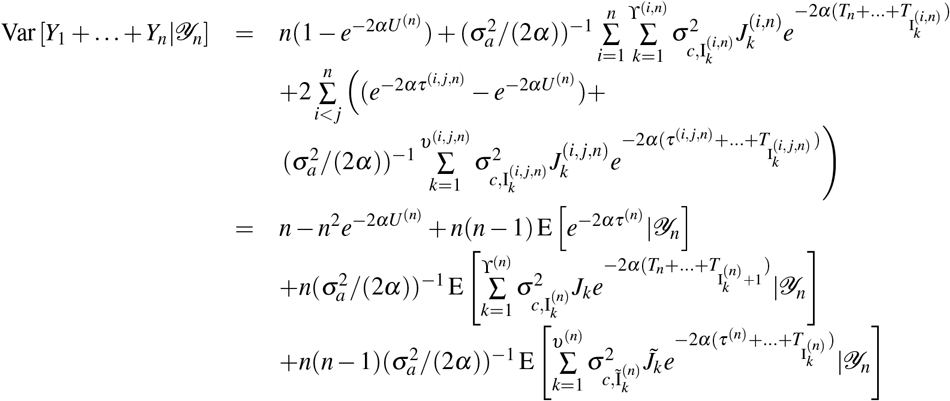

This immediately entails the second moment.

### Lemma 5.2

*Assume that the jump probability is constant, equalling p, at every speciation event. Let*

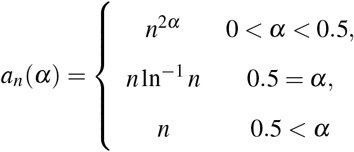

*and then for all α >* 0 *and n greater than some n*(*α*)

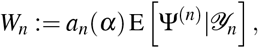

*is a submartingale and furthermore W_n_ converges a.s. and in L*^1^ *to a random variable W*_∞_ *with expectation*

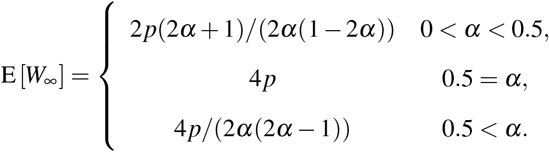

### PROOF PROOF

FOR *α >* 0.5

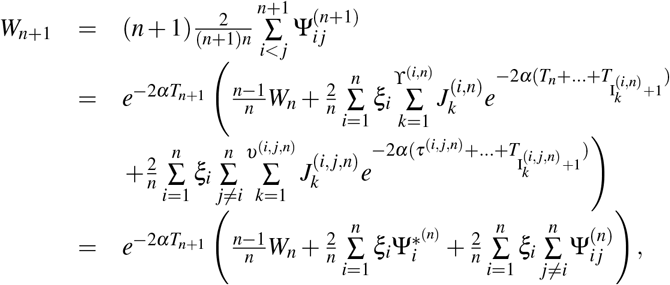

where *ξ_i_* is a binary random variable indicating whether it is the *i*–th lineage that split (see Fig 5).

**Figure 5:**
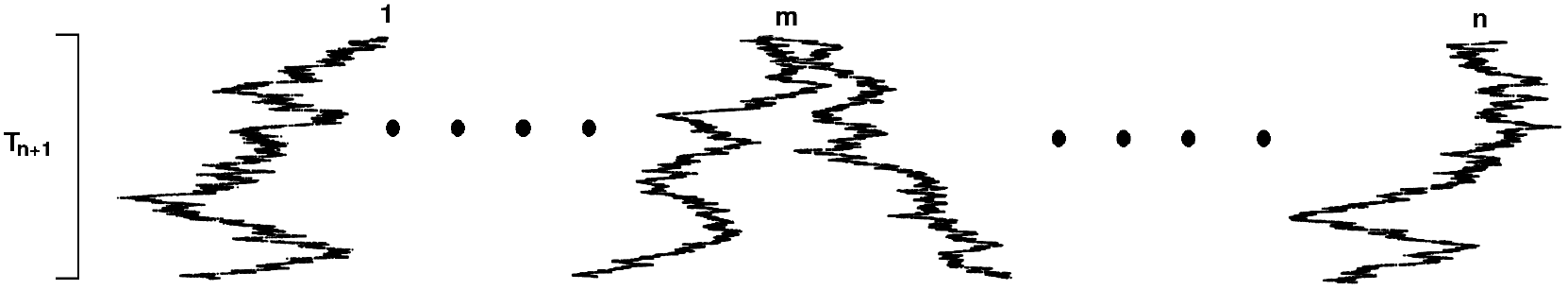
The situation of the process between the *n*–th and *n* + 1–st split. Node *m* split so *ξ_m_* = 1 and *ξ_i_* = 0 for *i ≠ m*. The time between the splits is *T_n_*_+1_ *∼* exp(*n* + 1).

Obviously the distribution of the vector (*ξ*_1_, ⋯, *ξ_n_*) is uniform on the *n*–element set {(1, 0, ⋯, 0), ⋯, (0, ⋯, 0, 1)}. In particular note

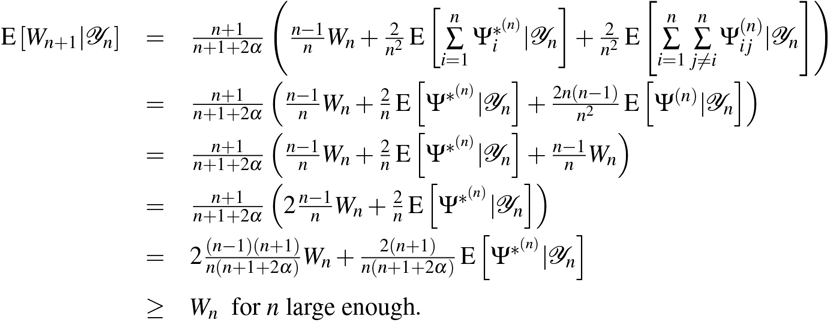

Therefore *W_n_* will be a submartingale with respect to *𝒴_n_* from a certain *n*. We know that E [*W_n_*] *< C_E_* for some constant *C_E_*, as E [*W_n_*] → 4*p/*(2*α*(2*α* − 1)) [Appendix A.2. 5] and hence by the martingale convergence theorem *W_n_ → W*_∞_ a.s. for some random variable *W*_∞_. As all expectations are finite E [*W*_∞_] *<* ∞. Furthermore Var [*W_n_*] *< C_V_*, for some constant *C_V_*, as, by Lemma 4.5, Var [*W_n_*] → 8*p*(1 *− p*)*/*2*α*, indicating E [*W_n_*] → E [*W*_∞_] and therefore E [*W*_∞_] = 4*p/*(2*α*(2*α* − 1)). Also the previous implies uniform integrability of {W_n_} and hence *L*^1^ convergence.

### PROOF

FOR *α* = 0.5 is the same as the proof for *α >* 0.5 except that we will have for *n* large enough

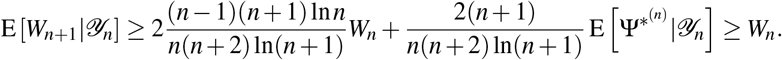

Now following again Bartoszek [5]’s Appendix A.2. we have E [*W_n_*] → 4*p* and Var [*W_n_*] → 8*p*(1 *− p*) (Lemma 4.5).

### PROOF

FOR 0 *< α <* 0.5 is the same as the proof for *α >* 0.5 except that we will have for *n* large enough

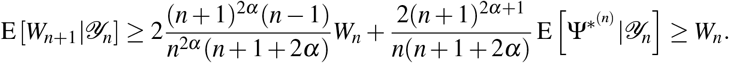

Following Bartoszek [5]’s Appendix A.2. we obtain E [*W_n_*] → 2*p*(2*α* + 1)*/*(1 − 2*α*) and Var [*W_n_*] converges to a constant by Lemma 4.5.

### PROOF OF THEOREM 3.1, PART (I), *α >* 0.5

We will show convergence in probability of the conditional mean and variance

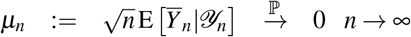

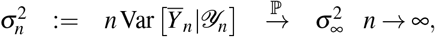

for a finite mean and variance random variable 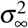. Then due to the conditional normality of *Y _n_* this will give the convergence of characteristic functions and the desired weak convergence, i.e.

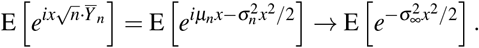

Using Lemma 5.1 and that the Laplace transform of the average coalescent time [Lemma 3 in 6] is

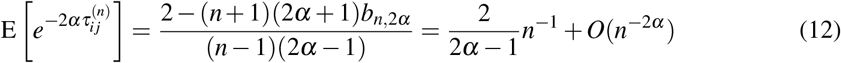

we can calculate

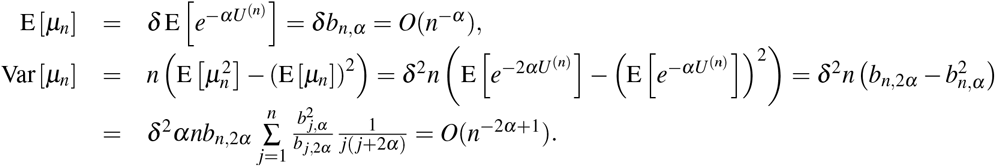

Therefore we have *µ_n_* → 0 in *L*^2^ and hence in P.

Lemma 5.1 states that

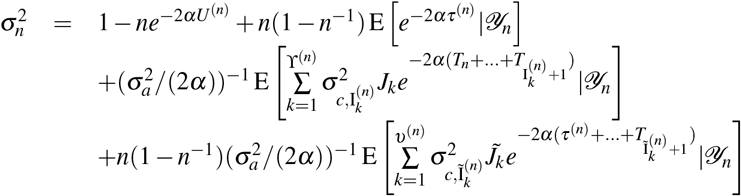

Using Lemmata 4.1, 4.3, 4.5, 5.2, Bartoszek [5]’s Appendix A.2 and remembering that *p_k_*, 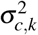 were assumed constant (equalling *p*, 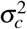 respectively), by looking at the individual components we can see that

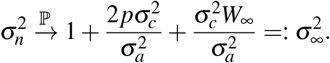

By Lemma 5.2

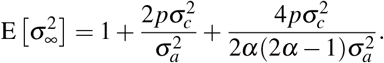

### PROOF OF PART (II), *α* = 0.5

This is proved in the same way as PART (I) except with the normalizing factor of the order of *n* ln^−1^ *n*.

### PROOF OF PART (III), 0 *< α <* 0.5

We notice that the martingale *H_n_* = (*n* + 1)*e*^(*α* +1)*U* (*n*)^*Y_n_* has uniformly bounded second moments. Namely by boundedness of *σ* ^2^, Lemma 5.1 and Cauchy–Schwarz

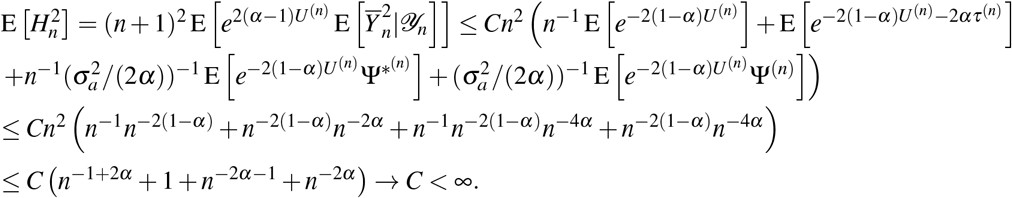

Hence, sup_*n*_ E 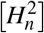 *<* ∞ and by the martingale convergence theorem, *H_n_ → H*_∞_ a.s. and in *L*^2^. As was done by Bartoszek and Sagitov [6] we obtain (Bartoszek and Sagitov [6]’s Lemma 9) *n^α^Y_n_* → *V* ^(*α−*1)^*H*_∞_ a.s. as in *L*^2^. Notice that for the convergence to hold in this regime, it is not required that 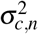 is constant, only bounded. We may also obtain directly the first two moments of *n^α^Y_n_* (however, for these formulae to hold, 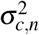 has to be constant)

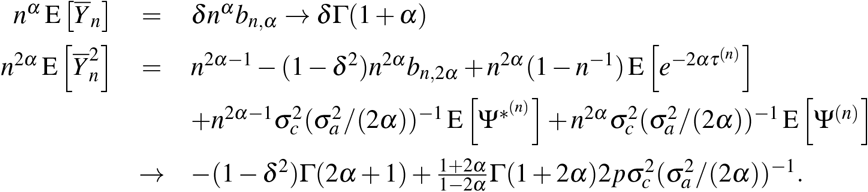

### PROOF OF THEOREM 3.5, PART (I), *α >* 0.5

From the proof of Part (I), Theorem 3.1 we know that *µ_n_* → 0 in probability. By the same ergodic argument as in Corollary 4.4 we obtain

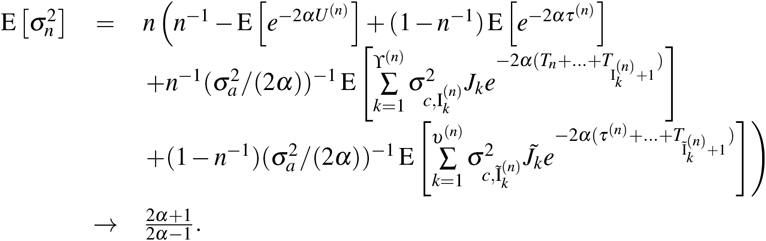

Furthermore Corollaries 4.4 and 4.6 imply

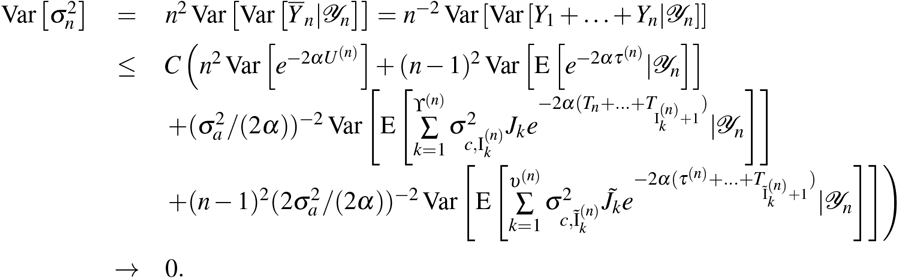

Therefore we obtain that 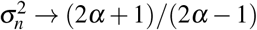 in probability and by convergence of characteristic functions

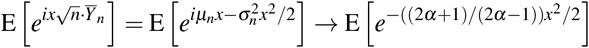

we obtain the asymptotic normality. Notice that on the other hand

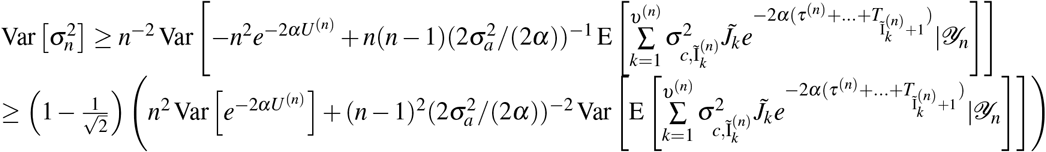

implying that the convergence 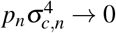 with density 1 is a necessary assumption for the asymptotic normality.

PROOF OF PART (II), *α* = 0.5 This is proved in the same way as PART (I) except with the normalizing factor of the order of *n* ln^−1^ *n*.

## Acknowledgements

KB would like to acknowledge Olle Nerman for his suggestion on adding jumps to the branching OU process, Wojciech Bartoszek for helpful suggestions concerning ergodic arguments, Venelin Mitov and Tanja Stadler for many discussions. KB was supported by the Knut and Alice Wallenberg Foundation.

